# Impact of an Oomen feeding with a neonicotinoid on daily activity and colony development of honeybees assessed with an AI based monitoring device

**DOI:** 10.1101/2020.02.04.933556

**Authors:** Gundula Gonsior, Frederic Tausch, Katharina Schmidt, Silvio Knäbe

## Abstract

Feeding experiments are standard tools in the pollinator risk assessment. The design (Oomen et al 1992) was developed to test insect growth regulators and herbicides. In recent years there was an update (Lückmann & Schmitzer 2015) on the outline in order to also focus on the advantage of different rates making a dose response design possible where exposure levels are known. Additionally, this design gives the possibility to test different rates for honey bee colonies foraging in the same landscape.

The main objective of the experiment presented here was to determine the natural variability of foragers losses of hives fed with a sub-lethal neonicotinoid concentration compared to an untreated control. Other objectives were to see if the neurotoxic exposure results in any observable sub-lethal effects and to find out if losses can be correlated to hive development. This was assessed with traditional methods and a novel, visual monitoring device.

## Introduction

In order to prove that a substance used in agriculture will bring no harm to pollinators, extensive testing must be performed on the active ingredients of plant protection products. There are several different testing protocols available. However, since there is a wide range of possible outside influences, tests run with free flying bees are always subject to uncertainty. One of the methods currently applied to compare bee mortality between different treatments is the use of dead bee traps. Regarding this method, potential uncertainties are known e.g. correlation of the total number of dead bees and the number of dead bees in the dead bee trap and the limited number of data sets which can be collected during testing. Furthermore, as the bees are foraging freely, it is very difficult to determine their level of exposure. Therefore, a realistic dose response design is not possible with spray application. The only test design, which gives the possibility to test different rates in the same environmental conditions, is the Oomen test design. The design presented was extended to include a digital hive monitoring device using computer vision and deep learning beside traditional mortality assessments. The device recorded all bees entering and leaving their hives with a camera, thus enabling the constant near-time observation of hive development and bee activity throughout the year. Deep learning analysis of the footage recorded made it possible to count the number of bees entering and leaving throughout the day and to calculate the losses of foragers over selected periods of time.

## Materials and Methods

To test the applicability of the approach, the study compared the hive development and losses of foragers from hives exposed to a neonicotinoid with a control group. Eight hives were monitored during the study. The colonies contained all brood stages (eggs, larvae and capped cells). Four colonies were fed with 500 g of sugar solution with a concentration of 200 µg imidacloprid/ kg of sugar solution over a period of ten consecutive days. The control group was fed the same amount of sugar solution during that period. Bees could fly freely, and had access to natural nectar and pollen sources. Each hive was equipped with a digital hive scale and an apic.ai monitoring system, consisting of a visual monitoring device and analysis software. It was attached to the hive entrance. It is solar-powered and UMTS-connected. All bees entering and leaving were recorded with a camera using infared light.

For further processing more than 4 TB of video footage was recorded and uploaded to a cloud. Concurrently, assessments of colonies, hive weight and daily mortality were made. The study was finished in September after a post-monitoring period of several months.

Presented were data of the first two brood cycles.

## Results

Mortality was assessed for each hive in the dead bee trap and a bottom drawer. For the first 28 days there was no clear increase in mortality detected, even during the feeding phase (Figure 1).

**Figure 1:**
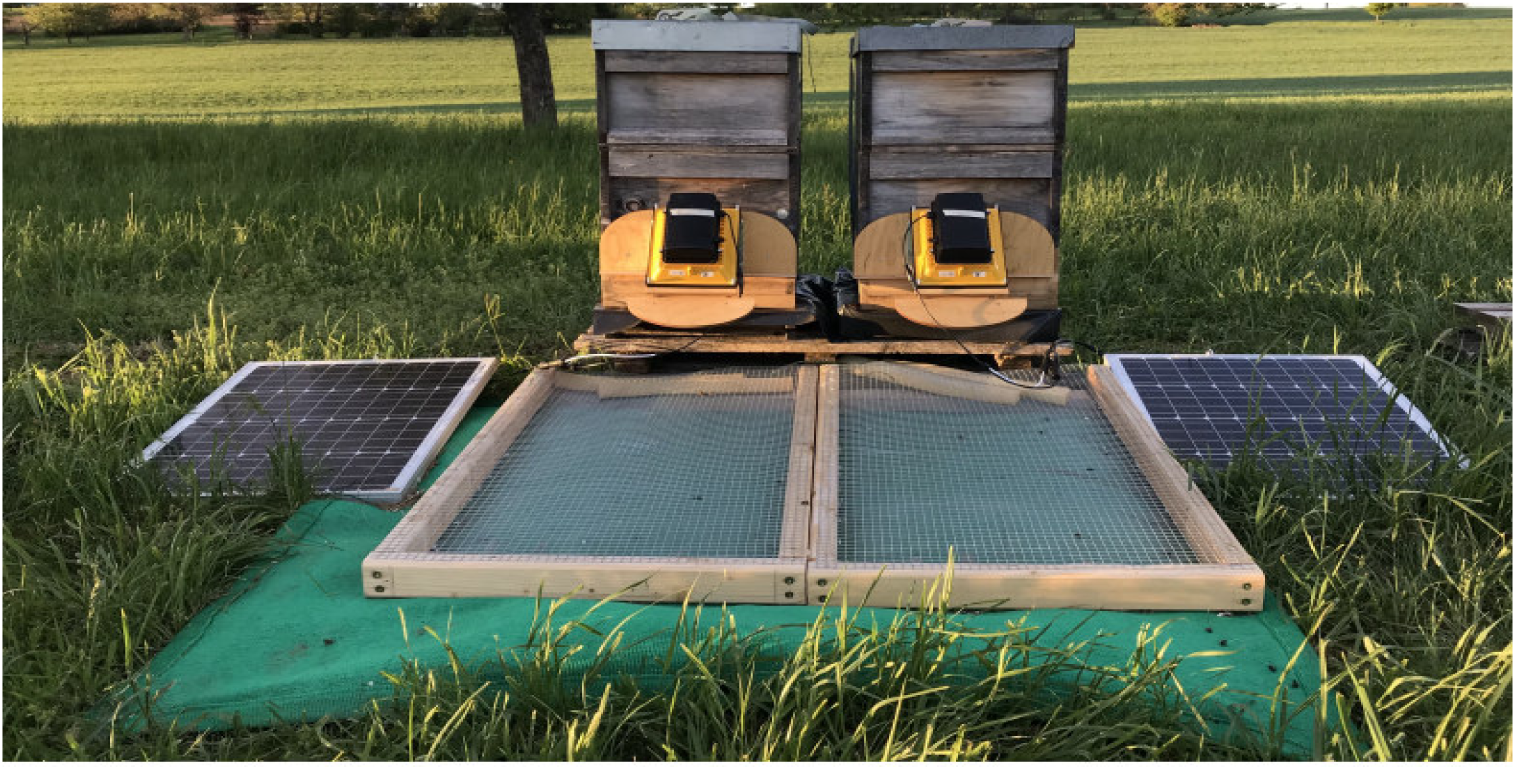
Hives with hardware system and bee monitoring.

Data variability is within an expected range.

Figure 4 shows effects on the strength of the colonies of the treatment group. A reduction in colony weight was expected since a neonic was fed in a concentration known to have effects. At the end of the 1st brood cycle a decrease in colony strength was observed in the treatment group. At the end of the 2nd brood cycle mean numbers were significantly lower compared to control data.

**Figure 2:**
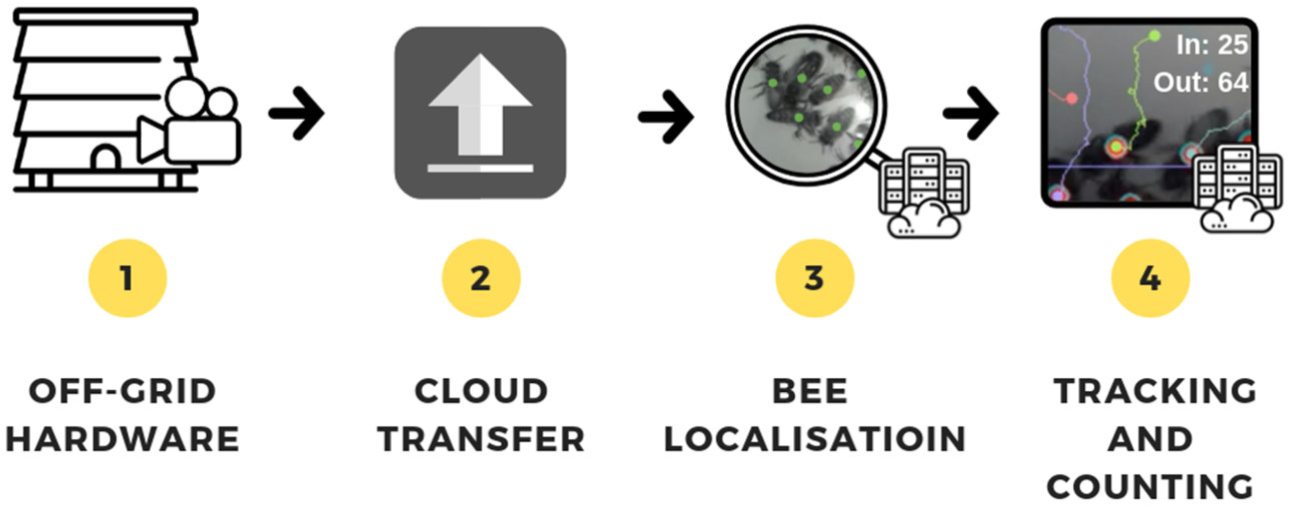
Operating principle of bee observation with digital monitoring device.

**Figure 3:**
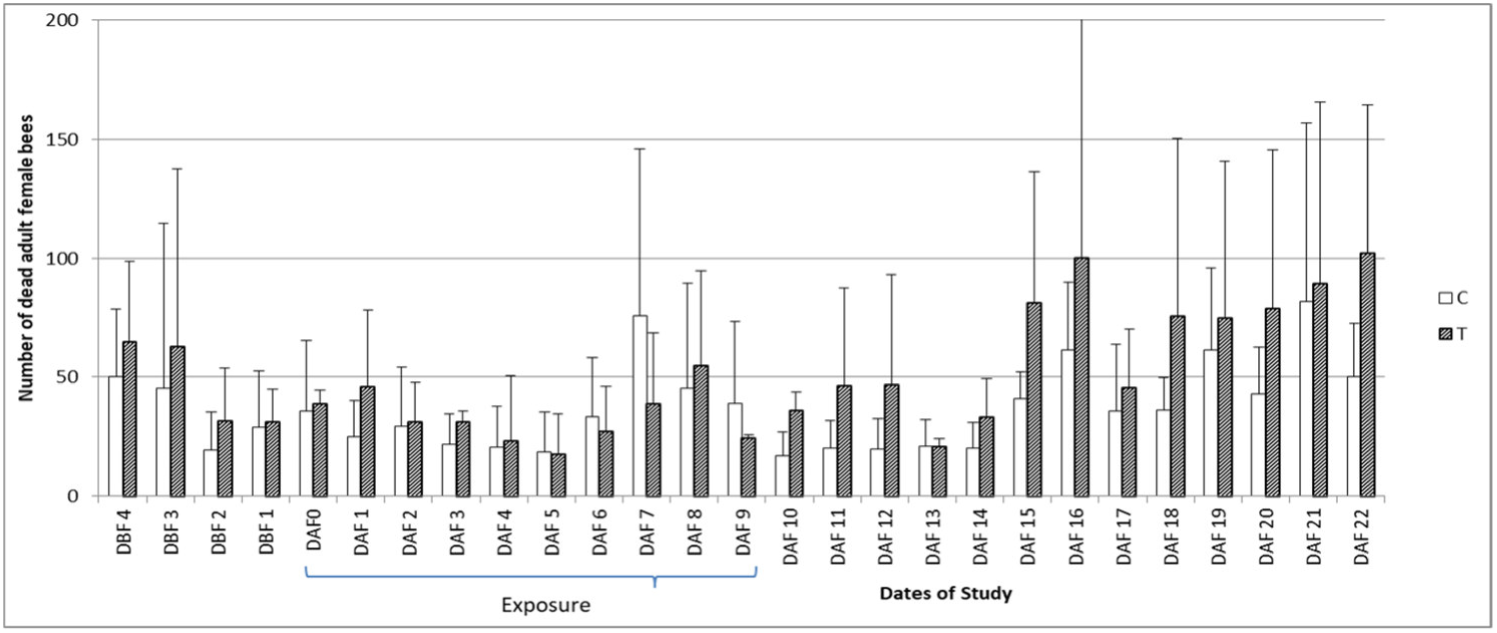
Mortality data assessed in dead bee traps and bottom drawer.

**Figure 4:**
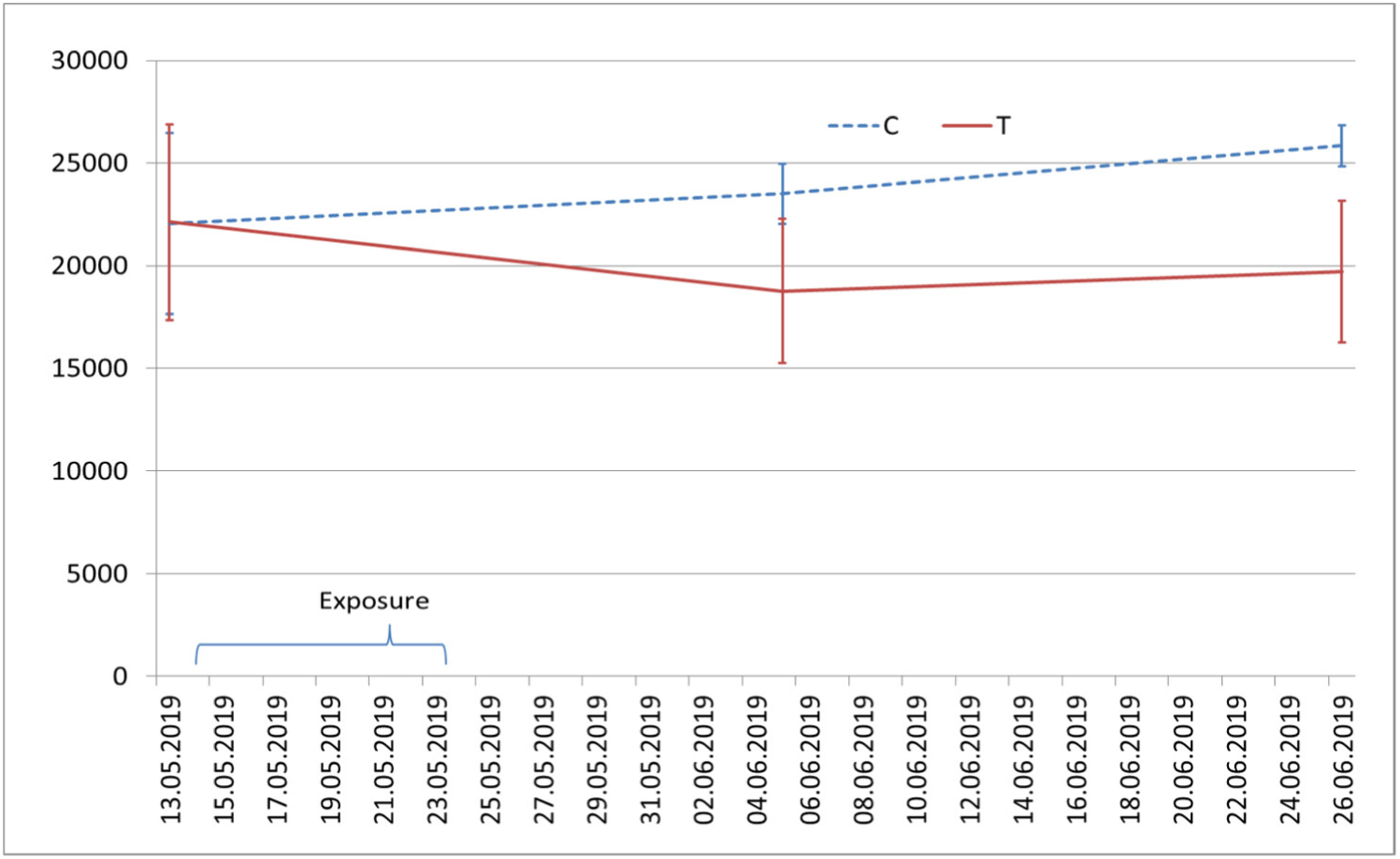
Development of colony strength over two brood cycles.

The difference in colony strength development correlated with the mean hive weights recorded (Figure 5). The mean of the treatment group followed the control, but never reached the same level.

**Figure 5:**
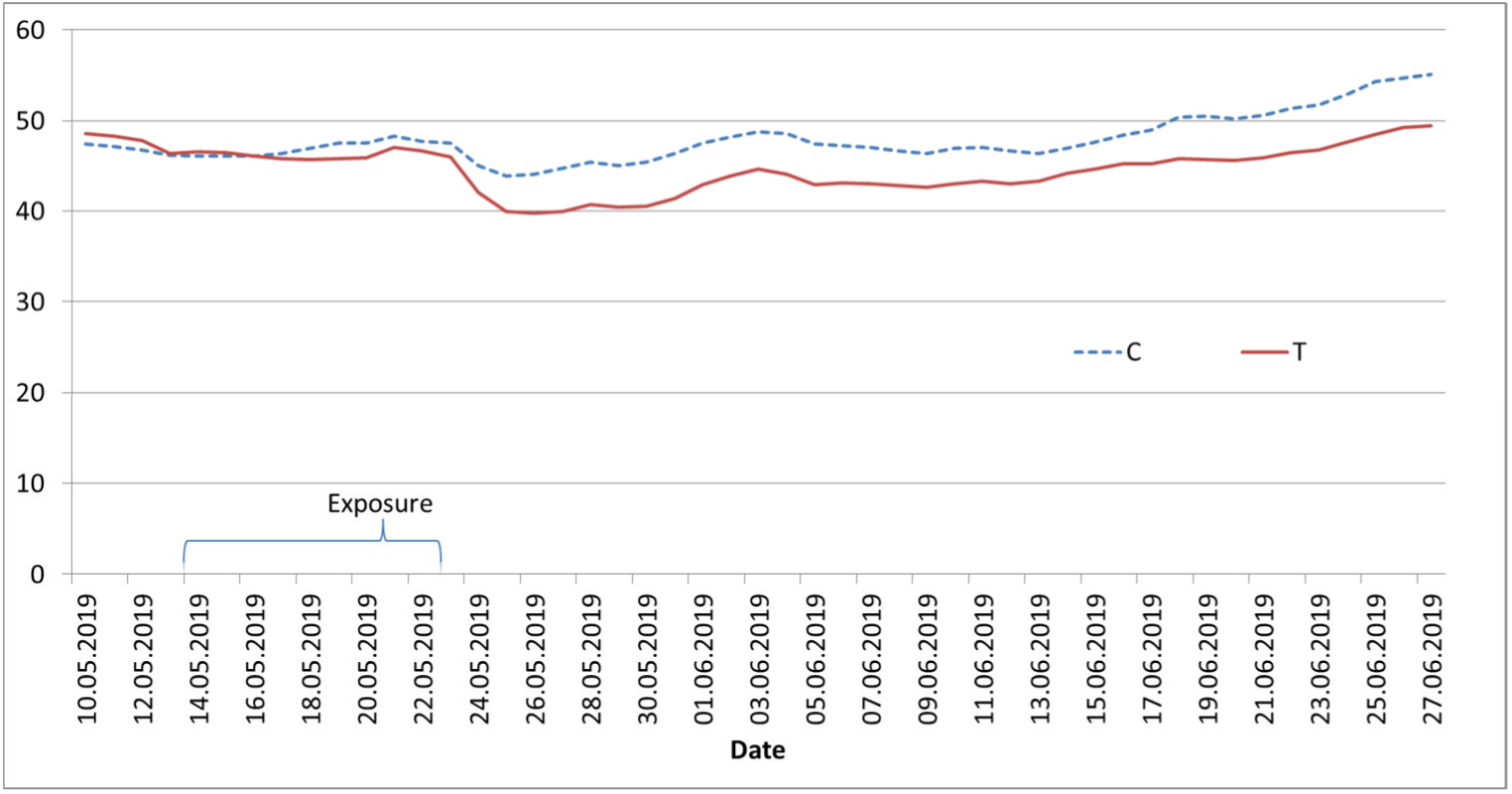
Mean weight of hives over two brood cycles.

Regarding the results of the apic ai monitoring, the following two figures show the activity pattern on different time scales. Figure 6 shows the change in activity per hive over the two brood cycles. Figure 7 presents activity in detail over the feeding period. The negative values in both figures represent the bees leaving the hive and positive values represent the returning bees. The Values presented are the sum of bees per hour.

**Figure 6:**
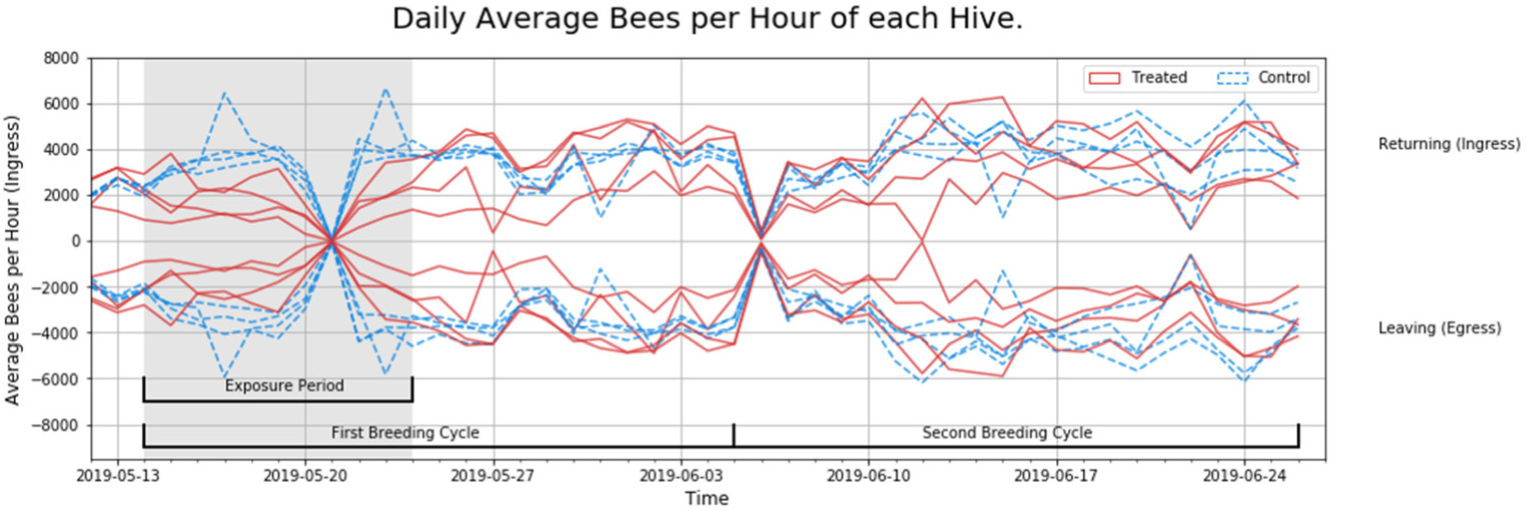
Daily average number of bees per hour over the two brood cycles.

**Figure 7:**
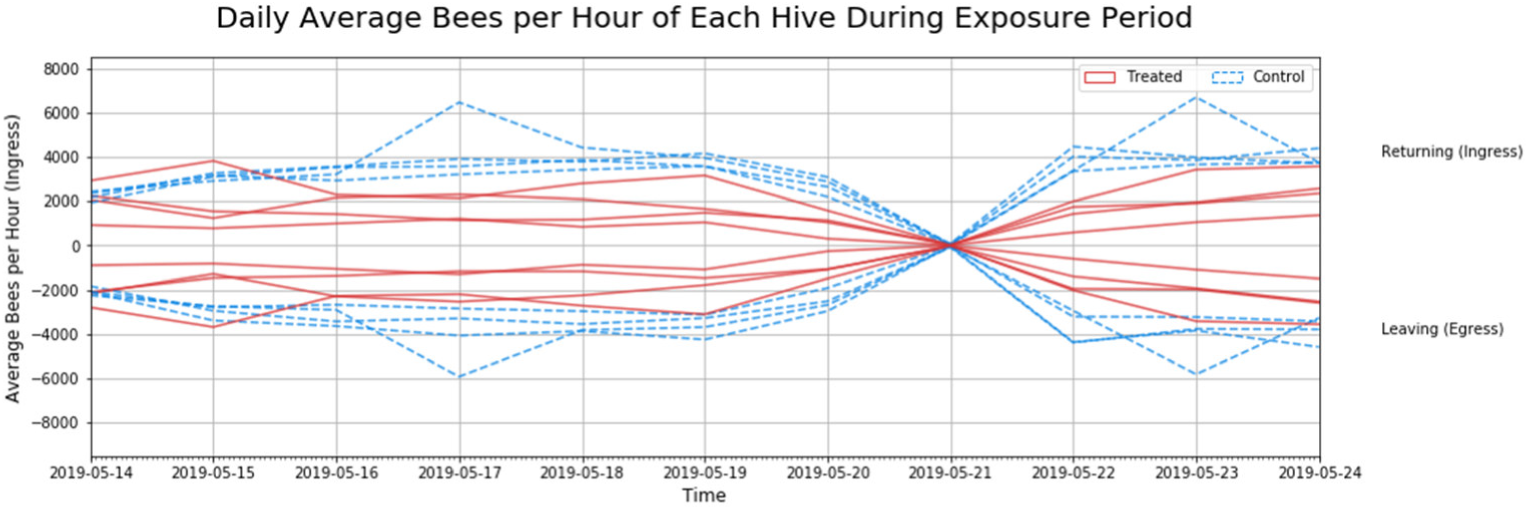
Daily average number of bees per hour over the feeding period.

During the exposure period (=feeding period) a higher activity was recorded for all of the control colonies starting from day 3 of exposure.

This is clearly visible in figure 7, which only covers the ten day feeding period.

At start of feeding activity was similar in treated and control colonies, afterwards activity decreased in the treatment group. On May 21^st^ 2019 inconvenient weather conditions resulted in a very low activity of all colonies. Results show when activity is reduced it has a straight influence on colony strength during the spring period. The most likely reason is the reduction in feeding of the larval feeding and consequently smaller amount of brood reared.

## Conclusions

AI based monitoring can help to explain effects due to more detailed observations. Sublethal effects on activity can be observed with the monitoring device. This is not possible with traditional methoology used in bee studies. The neurotoxic sub-lethal reductions correlate with reduction in colony strength and colony weight of the imidacloprid treated hives. However, even with 24 h observation data is variable like traditional methods. Further analysis will show what number of hives is needed in order to gain reliable data. Since constant determination of total loss of honeybees/hive is possible it would be advisable to include the methodology in regulatory studies. The method will give additional information not available at the moment. Nevertheless, traditional measurements are still needed to understand patterns observed in the field. Additionally the honey bee colony is the unit that has to be protected not the individual honey bee. Sub-lethal effects with no influence on the longterm vitality of the colony are interesting but not necessarily an obstacle to register a plant protection product.

One further advantage of the visual monitoring device is the fact that it enables a blind study analysis of the results obtained. That is possible because the data analysis of the activity can be done independent of the data collected in the field.

In future the visual monitoring device will also include pollen assesments and locomotion analysis of bees during their movment in and out of the hive.

## References

Lückmann J, Schmitzer S. The effects of fenoxycarb in a chronic Oomen feeding test – results of a ring test. Julius-Kühn-Archiv, 450, 2015

Oomen P.A., De Ruijter A. & J. Van Der Steen (1992): Method for honeybee brood feeding tests with insect growth-regulating insecticides. - EPPO Bulletin 22, 613 – 616.

